# Illuminating morphogen and patterning dynamics with optogenetic control of morphogen production

**DOI:** 10.1101/2024.06.11.598403

**Authors:** Dirk Benzinger, James Briscoe

## Abstract

Cells use dynamic spatial and temporal cues to instruct cell fate decisions during development. Morphogens are key examples, where the concentration and duration of morphogen exposure produce distinct cell fates that drive tissue patterning. Studying the dynamics of these processes has been challenging. Here, we establish an optogenetic system for morphogen production that enables the investigation of developmental patterning *in vitro*. Using a tunable light-inducible gene expression system, we generate long-range Shh gradients that pattern neural progenitors into spatially distinct progenitor domains mimicking the spatial arrangement of neural progenitors found in vivo during vertebrate neural tube development. With this system, we investigate how biochemical features of Shh and the presence of morphogen-interacting proteins affect the patterning length scale. We measure tissue clearance rates, revealing that Shh has an extracellular half-life of about 1h, and we probe how the level and duration of morphogen exposure govern the acquisition and maintenance of cell fates. The rate of Shh turnover is substantially faster than the downstream gene expression dynamics, indicating that the gradient is continually renewed during patterning. Together the optogenetic approach establishes a simple experimental system for the quantitative interrogation of morphogen patterning. Controlling morphogen dynamics in a reproducible manner provides a framework to dissect the interplay between biochemical cues, the biophysics of gradient formation, and the transcriptional programmes underlying developmental patterning.

## Introduction

Morphogen gradients play a crucial role in the spatial patterning and regulation of developmental programmes across various tissues. The processes governing morphogen production, dissemination, signal transduction, and gene regulation are dynamic and regulated by complex feedback mechanisms that span a range of time and length scales [1]. Achieving a quantitative and dynamic understanding of how robust and precise patterning arises requires the ability to perturb and measure morphogen production and cellular responses with spatial and temporal precision. This has been challenging in living embryos. Recently developed stem cell differentiation models hold promise for dissecting this problem *in vitro*, given their greater amenability to genetic manipulation, chemical exposure and imaging, but current models typically lack or are highly variable in their spatial organisation [2]. This limits quantitative studies. Consequently, there is a need for bespoke *in vitro* systems to study developmental patterning from morphogen gradient formation to cellular decision-making, which allow the precise control of morphogen dynamics in a reproducible tissue geometry.

Although methods that combine morphogen producing cells with non-producing receiving cells have been developed [3,4], these offer limited control over the positioning and extent of the source of morphogen producing cells. Light-based (optogenetic) regulation of protein function has emerged as a promising alternative for superior spatiotemporal control of gene expression [5]. Optogenetic control of morphogen signalling has revealed temporal requirements for cell fate determination and enabled the dissection of developmental gene expression programs [6–9]. Recent work has shown that spatial regulation of Sonic hedgehog (Shh) production, using optogenetic Cre recombinase systems, can generate signalling centres and patterning in stem cell differentiation systems [10,11]. However, although these systems provide control over the location and timing of morphogen production, they are irreversible and result in the continuous production of morphogen once initiated. A system that allows full spatial and temporal control of morphogen production is needed.

A technical challenge in developing optogenetic systems is to engineer a suitable dynamic range. Many systems capable of high levels of activation exhibit leakiness with low-level activation even without light stimulation. Conversely, tightly-regulated systems frequently activate only to modest levels. We sought to develop a system that offers stringent spatiotemporal regulation over both activation and inactivation of morphogen production. To this end, we focused on Shh-mediated patterning of the ventral neural tube, where Shh secreted from the notochord and floor plate forms a ventral-to-dorsal gradient specifying distinct progenitor domains along the dorsoventral axis (**Fig. 1A**).

**Figure 1.**
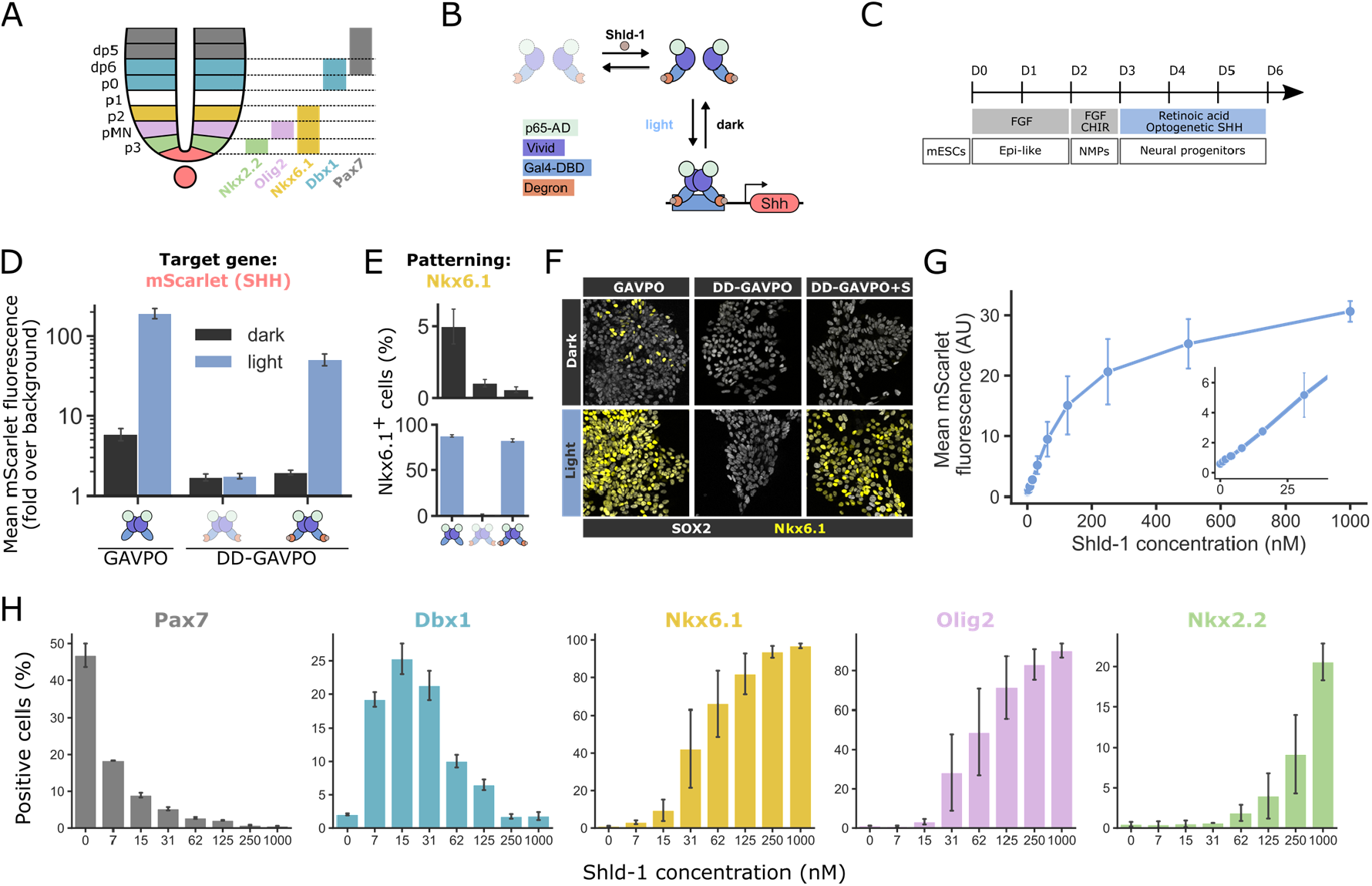
Optogenetic control of Shh expression and ventral neural progenitor identity using an optimised light-responsive transcription factor. **(A)** Schematic of progenitor domains and gene expression in the ventral neural tube. **(B)** Schematic of the optimised, optogenetic gene expression system based on the TF GAVPO fused to a Shld-1 responsive degron. Shld-1 stabilises GAVPO, and blue light illumination mediates TF dimerization, target promoter binding, and transcription of Shh-IRES-mScarlet. **(C)** Schematic of *in vitro* differentiation of mESCs to neural progenitors. **(D-E)** Effect of blue-light illumination on mScarlet (D) and Nkx6.1 (E) expression at D6 of neural differentiation measured by flow cytometry. GAVPO (left) and DD-GAVPO without (middle) or with 1 μM Shld-1 (right). **(F)** Representative microscopy images of Sox2 and Nkx6.1 expression at D6 of differentiation in dark (top) and blue-light illumination (bottom). **(G)** Dose-response of mean DD-GAVPO target gene expression to different concentrations of Shld-1 under blue light illumination. Cells were analysed by flow cytometry at D6 of neural differentiation. **(H)** Effect of Shld-1 concentration on the expression of patterning markers (Pax7, Dbx1, Nkx6.1, Olig2, and Nkx2.2, left to right). Cells were illuminated with blue light from D3 and marker expression was quantified at D6 by flow cytometry. Data and error bars of all experiments represent the mean and SEM of four (D, E) or two (G, H) independent experiments.

To enable tight, optogenetic regulation of Shh production, we developed an optimised expression system based on dual small-molecule and light control. Engineering this system into an *in vitro* stem cell model of neural tissue demonstrated tunable levels of gene expression that mimicked morphogen-like graded responses. Micropatterned stem cell colonies with localised Shh production recapitulated the long-range gradients and progenitor domains characteristic of neural tube patterning. We investigated how biochemical features of Shh and the presence of an extracellular morphogen-interacting protein affects the size of progenitor domains. We also used the system to measure the rate of Shh clearance, which gave new insight into the temporal aspects of gradient formation. Finally, we show how the level and duration of morphogen exposure dictate the acquisition and maintenance of positional identity. Overall, the ability to precisely manipulate morphogen dynamics represents a powerful system to model developmental patterning and dissect the molecular and cellular basis for morphogen gradient formation and interpretation.

## Results and Discussion

### Tight, optogenetic regulation by combined light and small-molecule control

To control the production of Shh during *in vitro* differentiation of mouse embryonic stem cells (mESCs), we used the optogenetic transcription factor (TF) GAVPO, which dimerises upon blue-light illumination resulting in target gene binding and transcriptional activation (**Fig. 1B**) [12]. We engineered mESCs to express GAVPO and a construct in which Shh together with a coexpressed fluorescent reporter (mScarlet) was driven by five GAL4/GAVPO binding sites in front of a synthetic YB-TATA minimal promoter, which provides low basal expression [13].

Following *in vitro* differentiation to neural tube progenitors [14], cells were illuminated with blue-light from Day3 (D3) to D6 (**Fig. 1C**). This resulted in a ∼30-fold increase in mScarlet expression in Sox2 expressing neural progenitor cells (**Fig. 1D**). Illumination resulted in most cells expressing Nkx6.1, a Shh responsive gene that marks the ventral p3, pMN, and p2 domains in the neural tube [15] (**Fig. 1A, E, F, Supplementary Fig. 1**). This indicated that GAVPO induced Shh expression is sufficient to activate the expression of genes characteristic of the ventral neural tube *in vitro*. However, we observed a proportion of cells expressing Nkx6.1 even in the absence of illumination (**Fig. 1E, F**), indicating that leaky activity of the system was sufficient to elicit a ventralising Shh signalling response.

Two major sources of undesired activity in optogenetic systems are induction by unintended/low-level illumination (resulting from routine handling of cells) and the inherent dark-state activity of many engineered light-responsive proteins [16,17]. The ability to limit GAVPO expression to a defined experimental time window would reduce mistimed activity. We reasoned that controlling GAVPO levels would enable us to work in an experimental regime that optimises the trade-off between dark-state activity-based leakiness and required maximal expression levels. To increase the controllability of the optogenetic system, we sought to regulate GAVPO expression itself. To achieve this, we fused GAVPO to a small-molecule (Shld-1) responsive conditional degron domain (DD-GAVPO, **Fig. 1B**) [18]. We found that the DD-GAVPO-based expression system showed minimal leaky expression in the dark (**Fig 1D**). In the absence of Shld-1, illumination did not induce mScarlet expression, consistent with GAVPO destabilisation. The addition of Shld-1 from D2 recovered high-fold-change blue-light induction of mScarlet (**Fig 1D**). Consistent with this, there was minimal Nkx6.1 expression in the dark, but Nkx6.1 induction after Shld-1 and light exposure was similar to the original system (**Fig 1E, F**). Thus, the optimised system lies at the sweet spot between low basal expression and robust inducibility.

### Tunable levels of DD-GAVPO activation with Shld-1

The ability to precisely tune the level of optogenetic activation is a desirable feature. We hypothesised that titrating the concentration of the small molecule Shld-1 could modulate the degree of DD-GAVPO activation, thereby offering fine control over optogenetic gene expression. The capability to precisely modulate activity in this way would represent a significant advantage over light input alone, as establishing illumination regimes that generate defined activity levels can be challenging. Quantification of light-induced mScarlet production across a range of Shld-1 concentrations revealed an approximately linear increase from 0-100 nM, with saturating activation achieved at ∼500 nM (**Fig. 1G**).

We investigated whether this tunable system recapitulated the graded response to Sonic hedgehog (Shh) signalling that directs the dorsal-ventral patterning of neural progenitors *in vivo* [18]. In the absence of Shld-1, light exposure failed to induce Shh expression, and cells expressed the dorsal marker Pax7. Strikingly, at low Shld-1 concentrations (<30 nM), optogenetic activation induced the intermediate neural tube marker Dbx1, characteristic of the dp6 and p0 domains. Moderate concentrations (>30 nM) allowed light triggered expression of the ventral markers Nkx6.1 and Olig2, specifying the pMN domain. High Shld-1 levels (>125 nM) allowed Nkx2.2 induction, a marker of the p3 domain (**Fig. 1A, H**). Thus, titrating Shld-1 concentration faithfully recapitulated the known relationship between Shh morphogen levels and neural progenitor identity, indicating that the conditional degron offers increased control over GAVPO activity.

In summary, the dual optogenetic and small molecule inputs of the GAVPO system synergize to enable tunable, tight regulation of gene expression output with minimal leakage. Taking advantage of the precise control of Shh expression in neural cells recapitulated the graded response to the Shh morphogen observed *in vivo*, a hallmark of morphogen signalling.

### Spatial regulation of Shh production *in vitro* recapitulates neural tube patterning

Next, we evaluated whether localised expression of Shh would result in long-range signalling activity and spatial patterning of *in vitro* neural progenitors. For this, we cultured differentiating ES cells on rectangular, laminin-coated micropatterns and illuminated one edge of each micropattern using a laser-scanning confocal microscope from D3 to D6 (**Fig. 2A**). Expression of mScarlet was restricted to cells in the illuminated area (**Fig. 2B, C**). Using two rounds of immunostaining, we evaluated domain-specific gene expression programmes ranging from the most ventral p3 to the intermediate dp6 domain (**Fig. 2B, C**). Strikingly, this revealed long range patterning. Nkx2.2, normally restricted to tissue close to the source of Shh in vivo, was expressed up to ∼15 μm beyond the illuminated domain. Olig2 and Nkx6.1 were expressed in a wider domain of ∼50 μm, with the expression of Nkx6.1 extending about one cell diameter further from the Shh source than Olig2. Dbx1, which is normally expressed in a stripe of neural progenitors in the middle of the neural tube, was expressed *in vitro* in a stripe of cells adjacent to the Nkx6.1 expressing domain. Thus, optogenetically regulated, localised Shh production *in vitro* recapitulates the spatial organisation of gene expression programmes characteristic of the neural tube (**Fig. 1A**).

**Figure 2.**
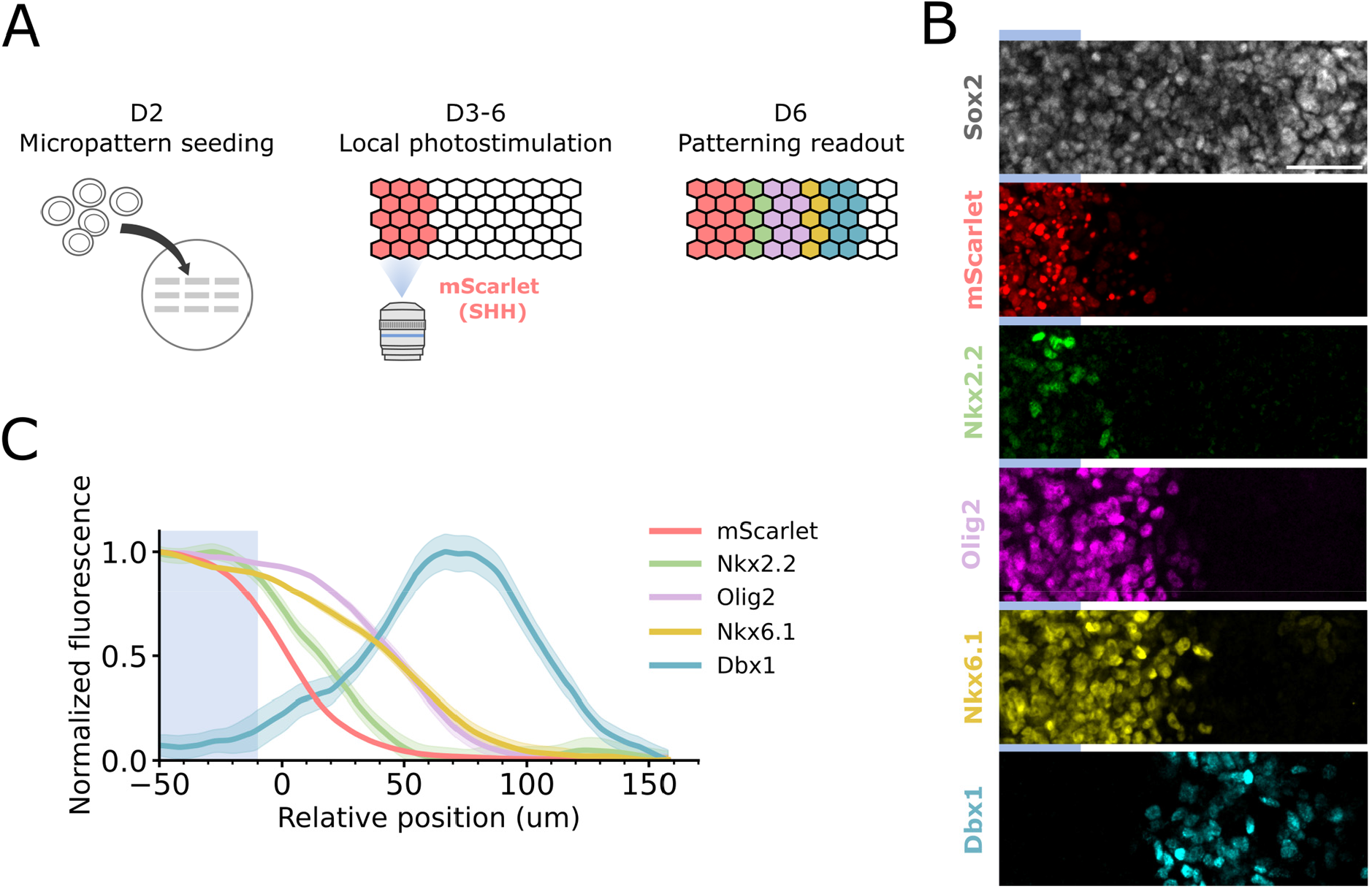
Spatially restricted Shh production recapitulates neural tube patterning. **(A)** Schematic of the protocol for spatially restricted Shh expression and patterning in micropatterned colonies. **(B)** Representative immunofluorescence images of patterning outcomes at D6. Images show a selected region of a 1500×300 μm rectangular micropattern. Localised Shh production was induced by illuminating a rectangular area with a width of 125 μm at the short edge of the micropattern (Methods). Blue bars above each image represent the spatial extent of illumination in the selected region. Scale bar = 50 μm. **(C)** Quantification of marker gene expression as a function of position, relative to half-maximal mScarlet fluorescence (Methods). Mean immunofluorescence signals for each marker were normalised by their maximal and minimal values. The mean (line) and SEM (shaded area) of gene expression from three independent experiments (with 4-6 patterns each) are shown.

The total extent of the optogenetically patterned region, defined as the distance from the limit of the mScarlet expression to the end of the Dbx1 expressing domain, was approximately 100 μM. This is similar to the *in vivo* Shh signalling length scale and the position of Dbx1 expressing cells in the mouse neural tube [20,21]. The sizes of the Nkx2.2 and Olig2/Nkx6.1 expressing regions *in vitro* were also comparable to those found in E9 mouse embryos [22]. To estimate the length-scale of the underlying Shh gradient, we made use of the Shld-1 titration experiments (**Fig. 1G, H**). Using mScarlet as a proxy for Shh levels and assuming an exponential gradient, we can construct a spatial map of marker gene expression that depends on the decay length of gradient (**Supplementary Fig. 2**). Comparing this mapping to the spatial pattern of gene expression resulting from localised Shh production (Fig. 2C) suggested a gradient decay length of ∼25 μm (**Supplementary Fig. 2**). This is consistent with the previously measured *in vivo* Shh gradient decay length of 20±5 μm [23]. The close correspondence between the structurally simple *in vitro* model and *in vivo* neural tube patterning was unexpected but suggests it offers a valuable tool for investigating aspects of morphogen gradient formation and progenitor fate acquisition.

### Progenitor domain size depends on biochemical properties of Shh

Having found that localised Shh production results in patterning with length scales comparable to *in vivo* neural tube development, we asked whether the patterning range was affected by biochemical properties of Shh. Mature Shh is C-terminally cholesterol-modified. This moiety has been implicated in controlling the spread of Shh through tissue and has raised the question of whether Shh remains membrane associated throughout its dissemination [24-26]. To investigate the effect of the Shh cholesterol modification, we optogenetically expressed two distinct forms of Shh: a version of Shh lacking the C-terminal cholesterol (NShh), and a version in which the transmembrane domain of CD4 had been appended to Shh in place of cholesterol (NShh-CD4) [27]. Cell-tethering via CD4 retained Shh’s ability to instruct progenitor ventralisation but abolished long-range signalling (**Fig. 3A, B**). This suggests that intercellular transmission of Shh normally involves Shh leaving expressing cells. By contrast, expression of NShh resulted in a 3-fold extension of the domain of Nkx6.1 expression, compared to Shh. This is consistent with a role for cholesterol in restricting Shh spread (**Fig. 3A, B**). NShh expression also resulted in an increase in the region of mixed Nkx6.1 and Dbx1 expression, hinting at the importance of controlled morphogen spread for establishing sharp gene-expression boundaries (**Fig. 3A**). These data demonstrate the requirement for Shh release from producing cells for optogenetic long-range patterning and illustrate how the biochemical properties of Shh contribute to the similarities of *in vivo* and *in vitro* patterning length scales.

**Figure 3.**
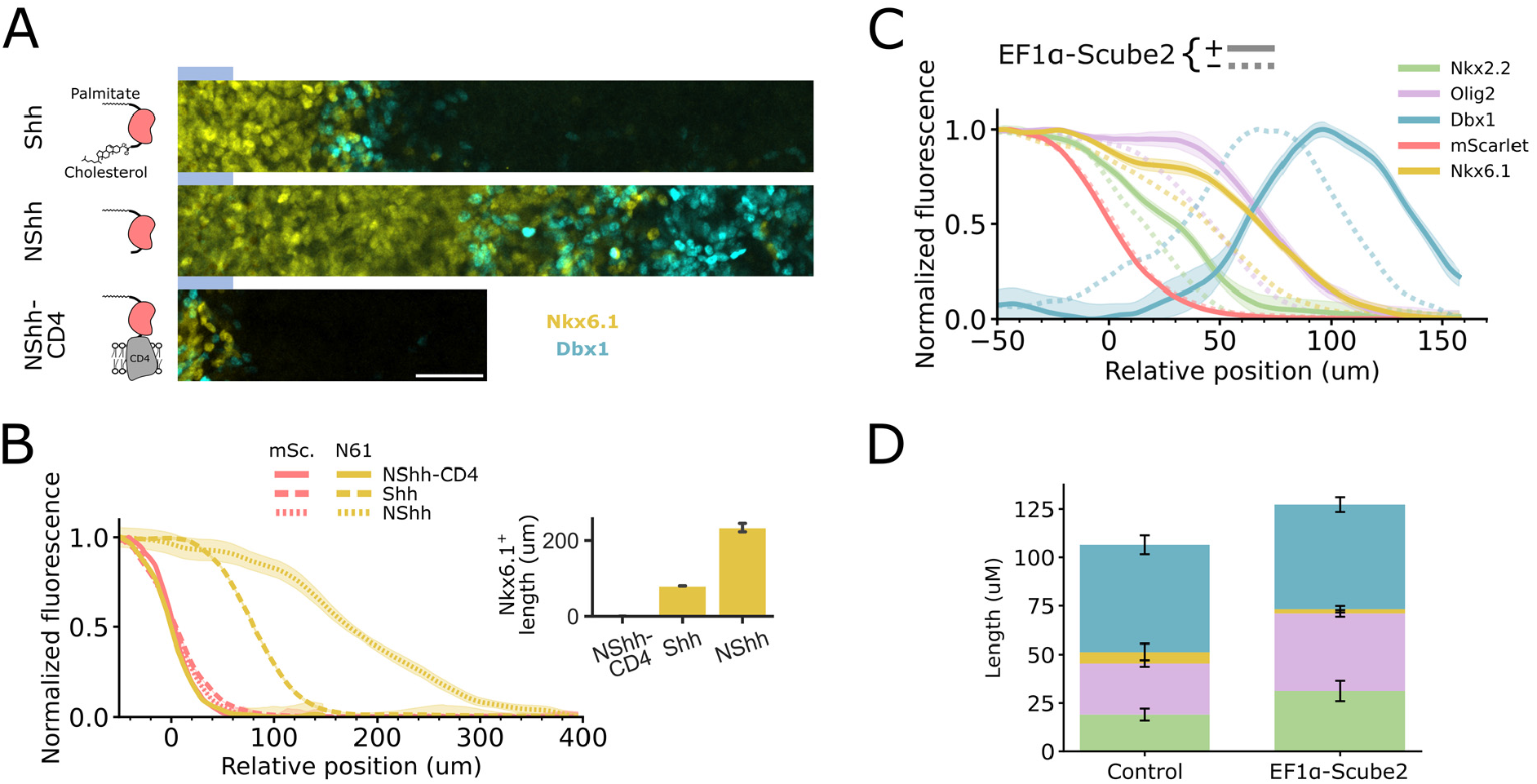
Modulation of gradient length scale by morphogen properties and extracellular binding partners. **(A)** Representative images of gene expression domains resulting from the optogenetic expression of different Shh variants at D6 of differentiation. Blue bars above each image represent the spatial extent of illumination. Scale bar = 50 μm **(B)** Quantification of Nkx6.1 (N61) expression as a function of position, relative to half-maximal mScarlet (mSc) fluorescence for different Shh variants. Mean immunofluorescence signals for each channel were normalised by their maximal and minimal values. The mean (line) and SEM (shaded area) of patterns from two independent experiments (with 4-6 patterns each) are shown. The bar plot shows the average length of the Nkx6.1^+^ domain (mean and SEM of two independent experiments). **(C)** Effect of heterologous Scube2 expression on patterning outcomes at D6 of optogenetic in vitro differentiation. Quantification of gene expression as a function of position, relative to half-maximal mScarlet fluorescence. Solid lines, ectopic Scube2 expression; dotted lines, wildtype (see Fig. 2C). **(D)** Comparison of gene expression domain lengths between heterologous Scube2 expressing and control cultures (Fig.2C, Methods).

### Scube2 expression increases patterning length-scale

In addition to their biochemical properties, the activity and spread of morphogens are influenced by interactions with extracellular binding factors. In the case of Shh, secreted SCUBE (Signal peptide, CUB, and EGF-like domain-containing) proteins cooperate with Disp1 to facilitate Shh release [28,29] and increase the range of Shh activity [30–32]. Moreover, in zebrafish a feedback loop between Shh signalling and Scube2 has been implicated in scaling the Shh gradient to the size of the neural tube [31]. However, how Scube2 expression affects mammalian neural tube patterning is not well understood.

To evaluate whether Scube2 expression affects progenitor domain size *in vitro*, we ectopically expressed Scube2 during optogenetic patterning. Strikingly, this resulted in an increased length of the ventral Nkx2.2 and Olig2/Nkx6.1 expressing domains by ∼60% and ∼40%, respectively (**Fig. 3C, D**). These data are consistent with the proposed function of Scube2 as a gradient extender in zebrafish [31] and illustrate how the optogenetic *in vitro* system can be used to study the modulation of morphogen signalling gradients.

### Dynamic regulation of Shh to measure clearance rate

Irrespective of molecular and cellular details, length and time scales of morphogen gradient formation are dictated by the effective transport and clearance rates of the morphogen through tissue [33]. Measurements of these rates have been achieved for multiple morphogens in Drosophila and zebrafish [34-36] but are largely lacking for mammalian systems.

Given the close resemblance of patterning length scales between the optogenetic *in vitro* system and the developing neural tube, we sought to use the system to measure the clearance rate of Shh. To do so, we optogenetically activated Shh expression for one day during in vitro differentiation, then transferred cells to the dark to halt Shh expression and measured extracellular and total Shh levels by immunofluorescence (**Fig. 4A**). Initially, total Shh staining resulted in a strong intracellular signal. The intracellular Shh signal disappeared within 4h of light withdrawal, indicating fast DD-GAVPO inactivation and Shh export/degradation. Extracellular Shh remained detectable after 4 h in the dark, but was completely cleared within 8h of light withdrawal.

**Figure 4.**
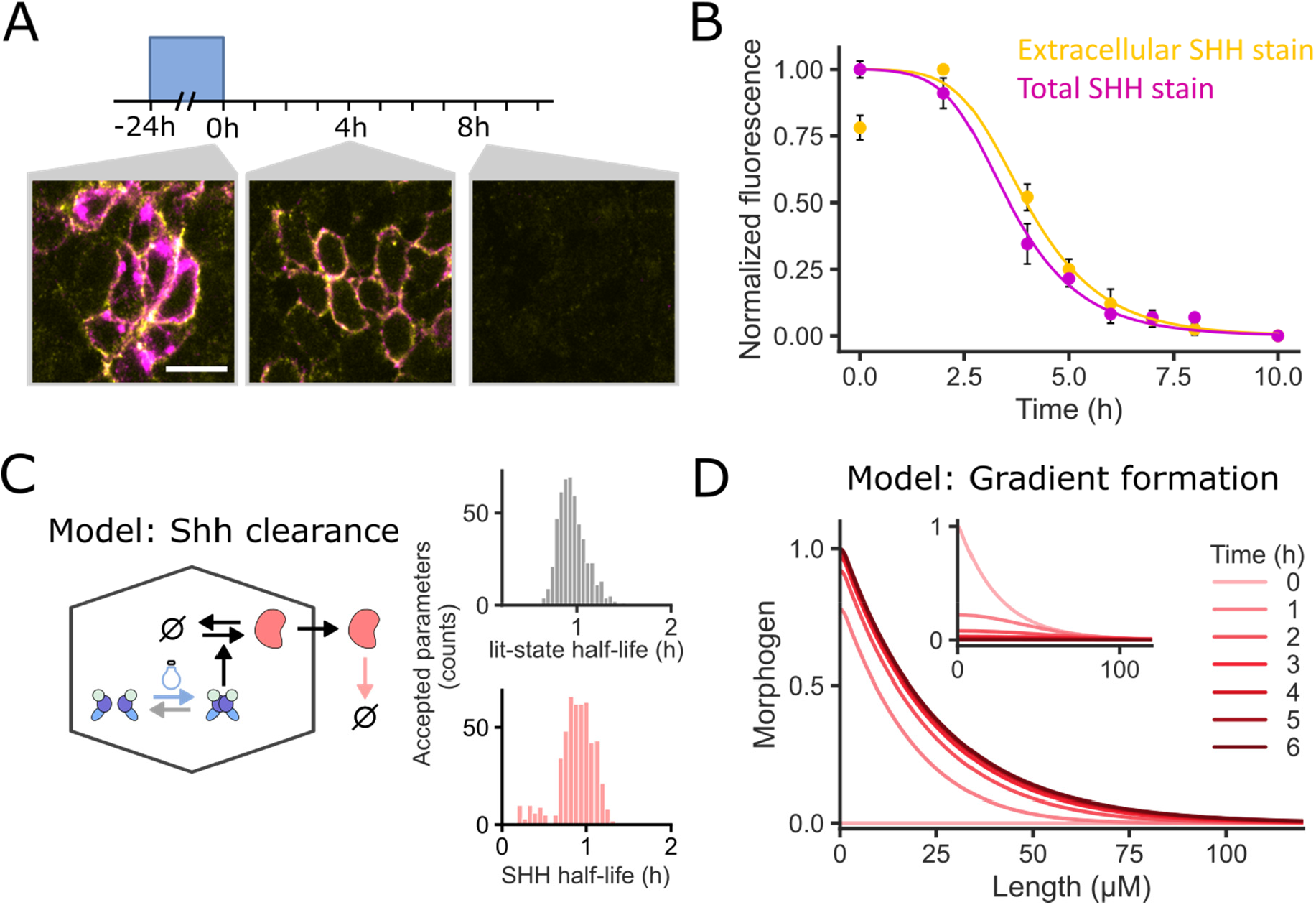
Quantification of Shh clearance. **(A)** Representative images of extracellular (yellow) and total (magenta) Shh immunofluorescence staining at 0, 4, and 8 h after blue-light withdrawal during in vitro differentiation. Cells were illuminated with blue light for 24 h before placing them in the dark. Scale bar = 10 μm. **(B)** Quantification of total (magenta) and extracellular (yellow) Shh immunofluorescence over time after blue-light withdrawal. Fluorescence values were normalised by their maximal and minimal values. Data points and error bars represent the mean and SEM of three independent experiments. Lines represent the averaged model simulation based on all 500 particles from the approximate posterior distribution (see (c)). **(C)** Model-based inference of DD-GAVPO dark-state reversion and Shh clearance rate. Schematic representation of a model for optogenetic Shh production, export, and clearance (left). Posterior parameter estimates for the DD-GAVPO lit-state (top) and Shh (bottom) half-life based on Approximate Bayesian Computation using the data shown in (B) (right, Methods). **(D)** Model of morphogen gradient formation upon activation of Shh production based on the Shh half-life estimate obtained in (c). Inset shows dynamics of gradient disappearance upon cessation of production.

The experimentally observed dynamics of total and extracellular Shh upon cessation of illumination depend on the rates of GAVPO deactivation as well as the export and degradation/clearance of Shh. To estimate these rates, we fit a simple mathematical model describing these dynamics to the experimental data (**Fig. 4B, C, Methods**). The experimental data constrained both the DD-GAVPO dark-state reversion and the extracellular Shh clearance rates (**Fig. 4C, Supplementary Fig. 3**). We estimated a half-life of the active DD-GAVPO lit-state to be ∼60min, extending the usefulness of the optimised expression system to applications that require highly dynamic gene expression perturbations. The half-life of extracellular Shh was estimated to lie between 0.5 and 1.5 h. This suggests that the clearance rate of Shh is in a similar range as Dpp in the *Drosophila* wing disc (∼ 45 min) [35] and Nodal in the zebrafish blastula (∼ 2h) [34].

Assuming a model of morphogen gradient formation in which a locally produced morphogen spreads and clears with constant rates throughout the tissue, we can calculate the effective morphogen diffusion rate (D = λ^2^ * k, where λ is the gradient decay length and k is the clearance rate). Assuming a Shh decay length of 25 μM (**Supplementary Fig. 2**) [23] and 0.22±0.05 × 10^-3^ s^-1^ for k (mean and STD of the posterior distribution corresponding to a half-life of 52.5 min, **Fig. 4C**), this results in an effective diffusion rate of 0.14±0.03 μm^2^/s, which lies in a similar range as Dpp and Wingless in the *Drosophila* wing disc [35]. This suggests that a steady state Shh gradient is reached within ∼3h from the start of morphogen production (**Fig. 4D**). This is faster than the time scales of the Shh signalling and downstream GRN response: patterning of mouse neural progenitors takes place over a period of 12-36h [37]. Thus, a Shh half-life of ∼1h suggests that the Shh gradient is continually renewed during neural tube patterning, making it amenable to temporal change (**Fig. 4D**).

### Progenitor identity is dynamically allocated

We next sought to use the optogenetic system to investigate the dynamics of progenitor identity determination. For this, we first compared patterning outcomes after one, two, and three days of light induction (**Figure 5A**). At D4, Nkx6.1 was strongly expressed in a domain with a length of ∼45 μm, while Olig2 was weakly expressed in a smaller region of progenitors. By D5, Dbx1 is expressed in a stripe adjacent to the Nkx6.1-expressing domain and Olig2 showed an expression pattern comparable to Nkx6.1, which remained stable at D6. Nkx2.2 started to be expressed in isolated cells at D5 and showed increased expression at D6. This progressive assignment of more ventral fates recapitulates progenitor identity determination *in vivo* [37,38]. The spatial extent of the Nkx6.1 expression domain was approximately constant from D4 to D6 (**Fig. 5B**). In agreement with the estimated rate of Shh clearance (**Fig. 4**), this result indicates that the length scale of the underlying Shh gradient reaches steady state before the downstream GRN. Thus, progressive ventralisation is not a consequence of the dynamics of gradient formation and is consistent with similar expression dynamics induced by a small-molecule agonist of Shh signalling *in vitro* [39,40].

**Figure 5.**
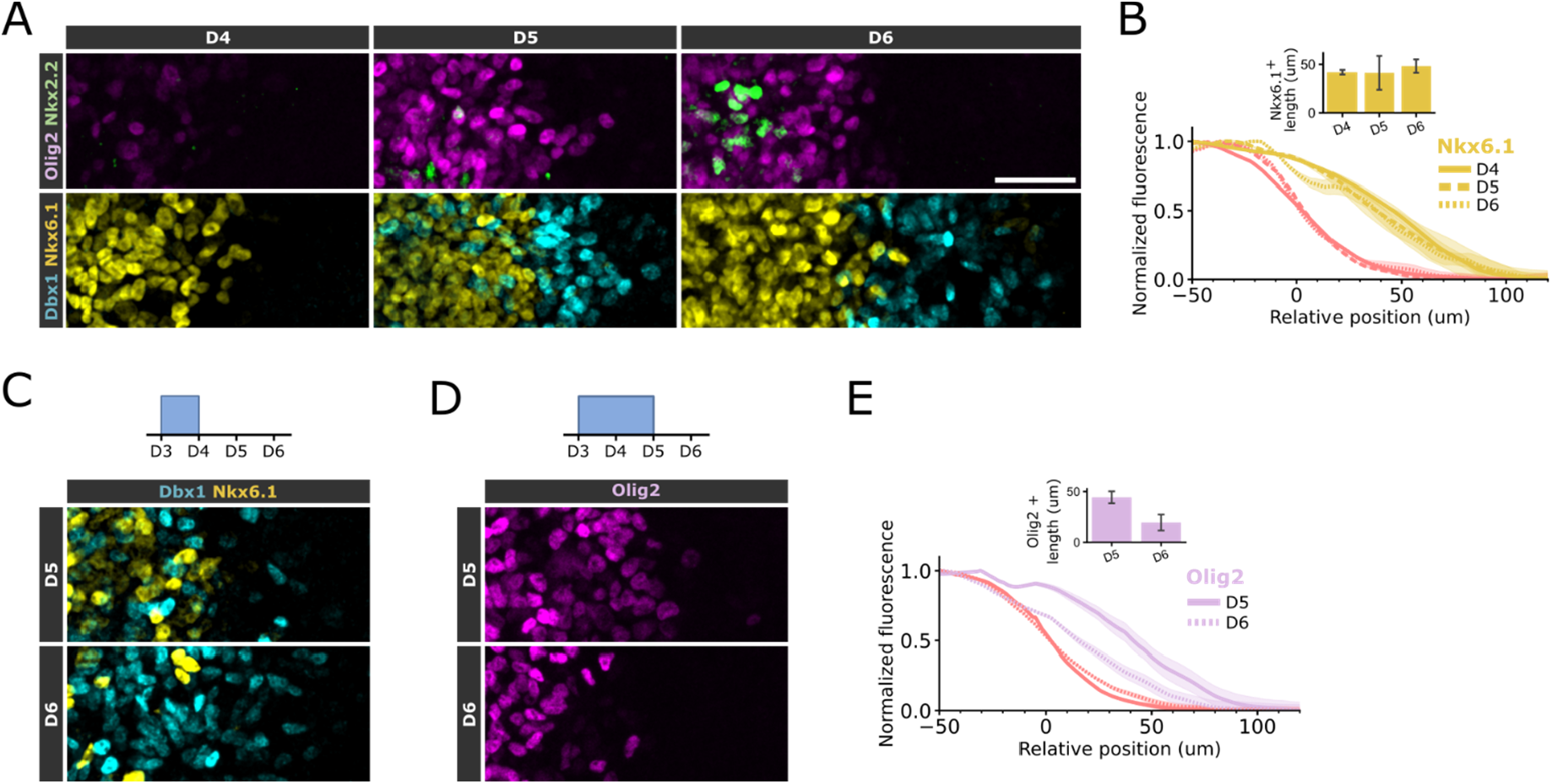
Dynamics of dorsal-ventral identity determination. **(A)** Representative microscopy images of patterning gene expression after one (left), two (middle), and three (right) days of localised Shh production. **(B)** Quantification of Nkx6.1 expression as a function of position, relative to half-maximal mScarlet fluorescence (red lines) at D4, D5, and D6. Mean immunofluorescence signals for each channel were normalised by their maximal and minimal values. The mean (line) and SEM (shaded area) of patterns from two independent experiments (with 4-6 patterns each) are shown. The bar plot shows the average length of the Nkx6.1 domain at the respective day of differentiation (mean and SEM of two independent experiments). **(C)** Representative microscopy images of Dbx1 (cyan) and Nkx6.1 (yellow) immunofluorescence at D5 and D6 in response to localised induction of Shh expression for 1 day. **(D)** Representative microscopy images of Olig2 immunofluorescence at D5 and D6 in response to localised induction of Shh expression for 2 days. **(E)** Quantification of Olig2 expression as a function of position, relative to half-maximal mScarlet fluorescence (red lines) at D5 (solid line) and D6 (dotted line) in response to two days of localised Shh induction (D). Mean immunofluorescence signals for each channel were normalised by their maximal and minimal values. The mean (line) and SEM (shaded area) of patterns from three independent experiments (with 4-6 patterns each) are shown. The bar plot shows the average length of the Olig2 domain at the respective days of differentiation (mean and SEM of two independent experiments). Scale bar = 50 μm.

Next, we used the ability to dynamically regulate Shh expression with DD-GAVPO (**Fig. 4**) to investigate maintenance of ventral identities. Following a one-day pulse of localised Shh production, light was withdrawn and Nkx6.1 expressing progenitors decreased over time, concomitant with an increase in Dbx1 expression (**Fig. 5C**), showing that continued Shh exposure is required for the maintenance of ventral progenitor identity [38]. However, a subpopulation of progenitors continued to express Nkx6.1 for at least two days after the cessation of Shh production, indicating the presence of hysteresis in ventral identity assignment [37]. Analysing the expression of Olig2 in response to a two-day pulse of Shh production further showed that cells closer to the Shh source maintained Olig2 expression for longer after light withdrawal than cells located further away (**Fig. 5D**). This resulted in a marked reduction in the length of the Olig2 expression domain after cessation of Shh production (**Fig. 5E**). Thus, the maintenance of progenitor identity depends on the level of past Shh exposure, conferred by cellular position in the gradient. In neural progenitors, Shh concentration affects both the level and duration of downstream signalling [38,41].

These experiments illustrate how spatiotemporal control over morphogen expression can be used to elucidate dynamic cellular decision-making during patterning. Together, these data show how the concentration and duration of Shh exposure contribute to both the determination and maintenance of ventral progenitor identity.

## Conclusion

By fusing an optogenetically regulated transcription factor to a small-molecule responsive conditional degron domain, we established an optogenetic system that enables precise spatiotemporal control over the production of the morphogen Sonic hedgehog (Shh) in an *in vitro* model of neural progenitor differentiation. The system provided low basal expression and robust inducibility. It recapitulated key features of embryonic neural tube patterning, including the formation of long-range Shh gradients and the specification of distinct ventral progenitor domains with appropriate spatial organisation.

We used the bespoke optogenetic morphogen system to dissect biochemical processes modulating Shh signal spread. Cholesterol modification of Shh and its extracellular binding partner, Scube2, affected the range of Shh signalling and consequently the scale of the patterned tissue. By dynamically controlling Shh production, we directly measured the clearance rate of Shh protein. The extracellular half-life of Shh protein of ∼1h suggests that Shh gradients are continually renewed during neural patterning. Finally, the system allowed us to probe the dynamics of neural progenitor fate acquisition and maintenance. We found that progressively more ventral fates are assigned successively, while the maintenance of a given ventral identity depends on both the level and duration of past Shh signalling experienced by a cell.

Overall, this optogenetic system establishes a novel experimental framework to measure the dynamic processes underlying morphogen gradient formation, interpretation, and transcriptional regulation during embryonic patterning. More generally, the availability of an optogenetic system that offers precise and tunable control of gene expression will be broadly applicable.

## Acknowledgements

We thank Tiago Rito for help with the micropatterning protocol, Despina Stamataki for testing of the Dbx1 antibody and Fei Chen, Patrick Hsu, and George Church for providing plasmids. We are grateful to Jake Cornwall Scoones, Anna Kicheva, David Willnow, and members of the Briscoe Lab for their constructive feedback on the manuscript. We thank the following Science Technology Platforms at the Francis Crick Institute for their expertise and assistance: Making Lab STP, Advanced Light Microscopy STP, and Flow Cytometry STP. This work was supported by the Francis Crick Institute which receives its core funding from Cancer Research UK (CC001051), the UK Medical Research Council (CC001051), and Wellcome (CC001051); and by Wellcome (226633/Z/22/Z). DB was supported by EMBO ALTF (671-2020).

## Author contributions

DB and JB conceived and designed the project, interpreted the data, and wrote the manuscript. DB performed experimental and computational analyses.

## Methods

### Cell culture of mouse embryonic stem cells

Mouse ESC lines were maintained in ESGRO Complete PLUS Clonal Grade Medium (Merck Millipore, Cat No. SF001-500P) supplemented with Penicillin/Streptomycin (Gibco, Cat No. 15140122) on tissue culture plates coated with 0.1% gelatin (Gibco, Cat no. G1393-100ML) at 37 °C with 5% carbon dioxide (CO_2_).

### Plasmid construction

All plasmids were constructed using golden gate assembly and propagated in *E. coli* TOP10 cells. We generated two piggyBac vectors with Esp3I cloning sites upstream of an EF1α-promoter-driven Puromycin (PBgg-Puro) and Hygromycin resistance gene (PBgg-Hygro), respectively (based on addgene #104536 [42]). To constitutively express GAVPO, we cloned its coding sequence followed by an SV40-terminator sequence downstream of an EF1α-promoter into PBgg-Puro (pDBB9). To generate DD-GAVPO (pDBB31), a sequence coding for the Shld-1 responsive degron [18] was synthesised by IDT and inserted upstream of the GAVPO coding sequence, separated by a glycine serine linker (GSGSGS). In order to control the expression of Shh light-responsively, we cloned the coding sequence of full-length mouse Shh behind a GAVPO-responsive promoter, consisting of five GAL4 binding sites and a synthetic YB-TATA minimal promoter into PBgg-Hygro (pDBB56). To measure GAVPO-mediated gene expression, an H2B-mScarlet-I fluorescent reporter [43] was expressed from the same transcript as Shh using the internal ribosome entry site (IRES2) of the *encephalomyocarditis* virus. Accordingly, we further cloned plasmids for the light-responsive expression of NShh (pDBB62) using the coding sequence for the first 197 amino acids of Shh and NShh-CD4 (pDBB63), adding the coding sequence of mouse CD4 downstream of NShh, separated by a glycine serine linker (GSGSGS). For the constitutive expression of Scube2, we generated a Cp36-recombinase donor plasmid based on pJT371 (addgene #184951) [44] containing the coding sequence of mouse Scube2 with a SV40-terminator sequence under control of an EF1α-promoter upstream of a Zeocin resistance cassette (pDBB57). Plasmid sequences were verified using nanopore sequencing (Full Circle Labs Ltd). All plasmids used in this study are summarised in **Supplementary Table 1**.

### Cell line generation

All cell lines are based on the mouse embryonic stem cell line HM1 [45]. PiggyBac and Cp36 integrations were performed by co-lipofecting the respective donor plasmids and a mouse codon-optimised piggyBac transposase [46] or Cp36 recombinase plasmid [44] in a 4:1 weight ratio using Lipofectamine™ Stem Transfection Reagent (ThermoFischer, Cat no. STEM00001). From 24 h after lipofection, cells were selected with either 1 μg/ml Puromycin, 100 μg/ml Hygromycin B, or 75 μg/ml Zeocin for at least 5 passages. Cell lines were generated by successive rounds of integration. To generate cell lines for the optogenetic regulation of full-length Shh, HM1 ESCs were first transposed with pDBB56 (UAS-Shh) and then pDBB9 (GAVPO) or pDBB31 (DD-GAVPO). For further characterisation experiments, we selected a clonal cell line for the DD-GAVPO-based expression of Shh by limiting dilution. To investigate the effect of Scube2 expression, this cell line was further modified by Cp36-mediated integration of pDBB57. For the comparison of Shh variants (Fig. 3A, B) wt ESCs were first transposed with pDBB31 (DD-GAVPO) followed by either pDBB56 (UAS-Shh), pDBB62 (UAS-NShh), or pDBB63 (UAS-NShh-CD4). All cell lines used in this study are summarised in **Supplementary Table 2**.

### Micropatterning of coverslips

Coverslips were micropatterned with rhLaminin-521 (ThermoFischer, Cat. No. A29249) as described in [47]: Coverslips were washed in isopropanol, activated in an UVO cleaner (Jelight) for 10 min, and incubated with 1 mg/ml PLL(20)-g[3.5]-PEG(5) (SuSoS) for 1 h at room temperature. After washing with ddH_2_O, the coverslips were placed on custom chrome photomasks (Compugraphics), ensuring close contact between mask and coverslip by pressing them using a plastic pipette tip. Coverslips were exposed to UV through the photomask for 8 min in the UVO cleaner, detached from the mask, incubated for 15 min in 70% ethanol, dried, and stored until usage for maximally 4 weeks. Coverslips were incubated with rhLaminin-521 diluted 1:10 in PBS +/+ (ThermoFischer, Cat. No. 14040091), at 37 C for 3h, washed 5 times in PBS +/+, and cells were seeded as described below.

### Neural progenitor differentiation

In vitro differentiation to neural progenitor cells [14] was performed in N2B27 media consisting of a 1:1 ratio of Advanced DMEM/F-12 (Gibco, Cat. No. 21331-020) and Neurobasal medium (Gibco, Cat. No. A35829-01), 1x N-2 and B-27 supplements, 2 mM GlutaMAX, 40 μg/ml BSA (Sigma-Aldrich, Cat No. A7979-50ML), and 0.1 mM 2-mercaptoethanol. At the start of differentiation (Day 0), mouse ESCs were plated onto Corning CellBIND tissue culture plates coated with 0.1% gelatin at a density of 6 × 10^3^ cells/cm^2^ in N2B27 media supplemented with 10 ng/ml bFGF (R&D, Cat. No. 100-18B). At day 2, the media was changed to N2B27 supplemented with 10 ng/ml bFGF and 5 μM CHIR99021 (Axon Medchem, Cat. No. 1386). If not otherwise stated, media was further supplemented with 1 μM Shld-1 (CliniSciences, Cat. No. AOB1848) for experiments with DD-GAVPO cell lines from Day 2 until the end of the experiment. From Day 3, media was supplemented with 100 nM RA (Sigma, Cat. No. R2625) with daily media changes. For global optogenetic Shh expression (experiments shown in Figs. 1 and 4), cells were transferred to an incubator equipped with a custom-built blue-light LED panel and illuminated every 5 min for 15 s at a light intensity of 400 μW/cm^2^.

For spatially localised optogenetic induction and measurement of Shh clearance, cells were seeded onto micropatterned coverslips on Day 2 of differentiation. For this, cells were washed with PBS, dissociated with Accutase (Gibco, Cat. No. 00-4555-56), centrifuged at 400g for 4 min, and resuspended in N2B27 supplemented with bFGF. 5 × 10^5^ cells were seeded per micropatterned coverslip. For spatial induction experiments, coverslips were secured in Attofluor cell chambers (ThermoFisher, Cat. No. A7816) and for Shh clearance experiments, coverslips were placed into 12-well tissue culture plates. Cells were allowed to attach for 2 h after which they were washed with N2B27 and media was changed to N2B27 supplemented with bFGF, CHIR99021, and Shld-1. Differentiation conditions before and after micropattern seeding are identical to those described above.

### Spatially localised illumination during differentiation

Spatial illumination of micropatterned cultures was performed using an inverted Olympus FV3000 laser-scanning confocal microscope with an Olympus UPlanSApo 10×-0.40NA objective, encased by a cellVivo environmental chamber at 37°C with 5% CO_2_. Micropatterns secured in Attofluor cell chambers were transferred to the microscope after media change to N2B27 supplemented with RA and Shld-1 at Day 3 of differentiation. To induce localised Shh production a rectangular area with a width of 125 μm at the short edge of rectangular micropatterns (1500 x 300 or 200 μm) was scanned every 10 min with a 488 nm laser at 1% laser power at 3.96x zoom and a scan speed of 2 μs per pixel (0.314 μm / pixel), resulting in a scan duration of approximately 1 s per pattern. This enabled us to control Shh expression in about 40 micropatterns per experiment. Media was changed daily. To measure the patterning response to Shh duration (**Fig. 5C-E**), illumination was stopped after 24 or 48 hours, and cells were left on the microscope until the end of the experiment/fixation.

### Immunofluorescence staining and imaging of differentiations

Cells were washed twice with PBS, fixed in 4% PFA for 10 min at room temperature, permeabilized in PBS/1% BSA/0.1% Triton-X100 (PBS-BSA-T) for 10 min, and incubated overnight at 4 °C with primary antibodies in PBS-BSA-T. Incubation with secondary antibodies was performed for 2 h at room temperature and imaging was performed in imaging buffer consisting of 700 mM N-acetyl-cysteine and 100 mM HEPES in ddH_2_O adjusted to pH 7.4.

To stain micropatterns for a full set of patterning markers, we performed two rounds of iterative immunofluorescence staining as described in the 4i protocol [48]. Briefly, after imaging, cells were washed with ddH_2_O and antibodies were stripped using three incubations in elution buffer (0.4 M L-Glycine, 2.3 M Guanidinium hydrochloride, 2.3 M Urea, and 54 mM Tris(2-carboxyethyl)phosphine hydrochloride in ddH_2_O adjusted to pH 2.5 with hydrochloric acid) for 10 min at room temperature. Cells were blocked in 4i blocking buffer (2% BSA and 150 mM Maleimide in PBS), washed with PBS-BSA-T, and the second round of immunofluorescence staining was performed as described above. In the first round, cells were stained with rat anti-Sox2 (1:1000, Invitrogen, Cat. No. 14-9811-80), goat anti-Olig2 (1:1000, R&D, Cat. No. AF2418), and mouse anti-Nkx2.2 (1:500, BD, Cat. No. 564731) primary antibodies and CS405 anti-rat (Biotium, Cat. No. 20419), AF647 anti-goat (Invitrogen, Cat. No. A21447), and AF488 anti-mouse donkey secondary antibodies (1:1000, Invitrogen, Cat. No. A21202). In the second round, cells were stained with rat anti-Sox2 (1:1000), mouse anti-Nkx6.1 (1:200, DSHB), and rabbit anti-Dbx1 (1:500, purified custom polyclonal antibody against the peptide antigen CDEDEEGEEDEEITVS generated by Cambridge Research Biochemicals, sequence provided by T. Jessell) primary antibodies and CS405 anti-rat, AF647 anti-mouse (Invitrogen, Cat. No. A31571), and AF488 anti-rabbit donkey secondary antibodies (1:1000, Invitrogen, Cat. No. A-31573).

Cells were imaged using an inverted Olympus FV3000 laser-scanning confocal microscope with an Olympus UPlanSApo 10×-0.40NA objective at 3.96x zoom, a 405 nm laser for CS405, 488 nm for AF488, 561 nm for mScarlet, and 640 nm for AF647. Z-stacks of 7 images spanning 23.28 μM were acquired. Scans for different fluorescence channels were performed sequentially to reduce spillover.

### Flow cytometry of fluorescent proteins and intracellular markers

1 ml/ml of LIVE/DEAD Fixable Dead Cell Stain Near-IR fluorescent reactive dye (ThermoScientific, Cat. No. L34976) was added to cell cultures and incubated for 30 min at 37 °C. Cells were washed twice with PBS, dissociated with Accutase, and collected by centrifugation for 4 min at 400 g in lobind Eppendorf tubes. Fixation in 4% PFA was performed for 10 min at room temperature and cells were washed in PBS and resuspended in PBS/0.5% BSA. Per experimental condition, 1 million cells were pelleted in V-bottom 96 well plates and incubated with primary or directly conjugated antibodies in PBS/0.5% BSA/TritonX-100 for 1.5 h at room temperature in the dark. If required, cells were incubated with fluorescently labelled secondary antibodies for 40 min at room temperature. After washing, cells were resuspended in PBS/0.5% BSA and analysed on a BD Fortessa analyzer (BD). Single cells were identified using forward and side scatter, a Sox2-positive progenitor population was gated for, and live cells were selected by gating for Dead Cell Stain negative cells (**Supplementary Fig. 1**). Throughout the paper, the mean of the fluorescence intensity distribution is presented for mScarlet and the percentages of positive cells are reported for patterning markers. The staining of patterning markers Fig. 1H was split into three different panels: rabbit anti-Dbx1 (1:500), mouse anti-Pax7 (1:500, DSHB), and rat anti-Sox2 (1:1000) with CS405 anti-rat, AF488 anti-rabbit, and AF647 anti-mouse goat secondary antibodies (1:1000); Nkx6.1-AF647 (1:100, BD Cat. No. 563338) and Sox2-V450 (1:100, BD Cat. No. 561610); goat anti-Olig2 (1:500), Nkx2.2-AF647 (1:100, BD Cat. No. 564729), and Sox2-V450 (1:100) with AF488 anti-goat secondary antibody (1:1000).

Quantification of mScarlet fluorescence in response to Shld-1 concentrations (Fig. 1G) was performed in live cells. For this, cells were washed twice with PBS, dissociated using Accutase, resuspended in PBS/5% foetal bovine serum (Pan Biotech, Cat. No. P30-2602)/0.1μg/ml DAPI, and kept on ice until analysis. Single cells were identified using forward and side scatter and live cells were enriched by gating for DAPI negative cells.

### Analysis of spatial gene expression patterns

Automated quantification of marker gene expression in micropatterned cultures was performed with custom Python scripts using the OpenCV and NumPy packages. For each image stack, a maximum intensity z-projection was performed. Micropatterns were automatically identified based on the Sox2 expression. First, the image was binarized using Otsu’s thresholding followed by morphological closing with a 20×20 kernel. The largest contour surrounding the pattern is then identified, approximated by a rectangle [49], and the image is cropped to remove areas outside the pattern.

For each channel of a pattern, the minimum image pixel intensity was subtracted from the image and the median absolute deviation of pixel intensities was used to identify and remove outlier pixels with user-defined thresholds. Average intensity profiles were generated along the axis perpendicular to the illuminated pattern edge. The profiles were smoothed using a LOWESS function (fraction = 0.075) and normalised to lie between 0 and 1 by subtracting the minimal intensity value and dividing by the maximal intensity value. To avoid variability resulting from potential differences in pattern alignment during illumination and imaging, intensity profiles of patterns were aligned to the point of half-maximal mScarlet fluorescence intensity for each experimental condition and repeat. Aligned curves were then averaged and normalised to lie between 0 and 1 to result in the profiles presented in this study. Lastly, domain sizes were calculated by defining the end of each domain as the point when their intensity profile drops below 20% of the maximal value and calculating the difference between adjacent boundaries. As such domains do not represent the exclusive expression of a single marker.

### Measurement of Shh clearance

To measure Shh clearance, we illuminated micropatterned colonies for 24 h from Day 3 to Day 4 of differentiation and performed extracellular Shh immunostaining in live cells at 0, 2, 4, 5, 6, 7, 8, and 10 h after transferring cells to the dark [50,51]. For this, we incubated cells with Shh (5E1) mouse antibody (1:500, DSHB) in N2B27 media containing 1% goat serum, supplemented with RA and Shld-1 for 40 min at 37 °C with 5% CO_2_. Cells were then washed once with PBS, fixed with 4% PFA for 20 min, and washed again two times with PBS. We next performed antigen retrieval by incubating cells in 1x citrate buffer (Agilent, Cat. No. S2369) for 45 min at 65 °C. Next, we performed a secondary antibody staining with AF488 goat anti-mouse (1:1000) in PBS/1% BSA for 2 h at room temperature. Cells were washed in PBS/1% BSA and then permeabilized by incubation in PBS-BSA-T for 10 min to enable total Shh staining. Cells were next incubated overnight at 4 °C with Shh (C9C5) rabbit antibody (1:500, Cell Signaling Technologies, Cat. No. 2207) in PBS-BSA-T, washed, and incubated for 2 h with an AF647 goat anti-mouse secondary antibody (1:1000) at room temperature. Coverslips were mounted using ProLong Gold antifade reagent (Invitrogen, Cat. No. P36930). Cells were imaged using an inverted Olympus FV3000 microscope with a Plan-Apochromat 20×-0.75NA objective. Z-stacks of 13 images spanning 24 μM were acquired in the centre of 6-8 micropatterns per condition with 3.71x zoom such that the scanned area did not extend over the micropattern border. Immunofluorescence intensity measurements were performed by calculating the mean pixel value of maximum intensity z-projected images that were background corrected by subtracting the 10th percentile pixel value. Per conditions, mean fluorescence intensities of micropatterns were averaged and values of a given time-series were normalised by subtracting the minimal intensity value and dividing by the maximal intensity value.

### Mathematical model and parameter estimation for Shh production and clearance

To estimate the rate of Shh clearance from experimental data, we set up a simple ordinary differential equation (ODE) model that describes GAVPO-based Shh production. The model consists of the following three ODEs describing light (I) dependant GAVPO (de-)activation (Eq. 1), GAVPO dependent production of cell internal Shh (Shh_int_) as well as its degradation and transport outside of the cell (Eq. 2), and extracellular Shh (Shh_ext_) clearance (Eq. 3):

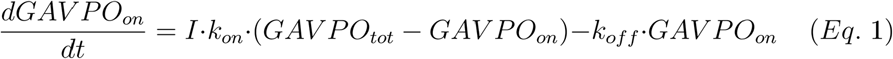

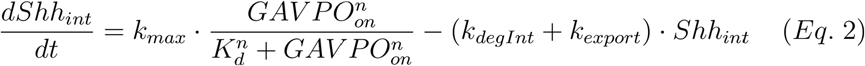

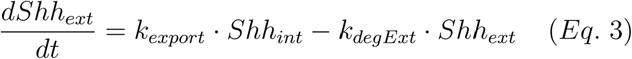

To infer parameter sets for which the ODE model describes well our experimental data (Fig. 4A, B), we used rejection approximate Bayesian computation (ABC) via the GpABC Julia package [52]. For ODE simulations, we set the initial conditions for GAVPO_on_ to 1. For each given parameter set, the initial conditions for Shh_int_ and Shh_ext_ were set to their steady-state values assuming GAVPO_on_ is fixed to 1. For ABC inference I and k_on_ were set to 0, given that blue-light illumination is stopped at the start of the experiment. k_max_ is set to 1 as it solely affects the absolute production levels which are not relevant for the parameter inference due to normalisation. The prior distributions of the remaining free parameters are summarised in **Supplementary Table 3**. Simulated data vectors for extracellular Shh staining (Shh_ext_) and total Shh staining (Shh_ext_+Shh_int_) were generated for the experimentally measured time points. Each vector was normalised through division by its maximal element. Parameter sets/particles were accepted if the Euclidean distance between the combined simulated data vector and experimental data vector was below a fixed threshold (0.25). ABC was stopped after 500 particles were accepted. Approximate marginal posterior distributions are summarised in **Fig. 4C** and **Supplementary Fig. 3**.

### Simulations of gradient formation and decay

To evaluate the dynamics of gradient formation expected from the experimentally measured Shh clearance rate (**Fig. 4 A-C**) and gradient length scale (**Supplementary Fig. 2**), we used a simple synthesis-diffusion-degradation model (Eq. 4), where S describes localised synthesis of Shh at the left edge of the domain at a constant rate, D is the diffusion rate of Shh (0.13 μM^2^/s) and k_deg_ is its clearance rate (0.22 × 10^-3^ s^-1^).

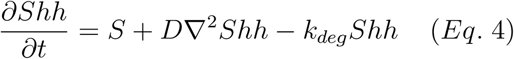

Simulations were performed using a finite difference scheme in Julia with the DiffEqOperators and DifferentialEquations packages [53] and reflective Neumann boundary conditions. Gradient formation was simulated on a 400 μM 1-dimensional domain discretized into 1000 segments. For gradient formation, the initial condition was Shh = 0 throughout the domain. To simulate gradient decay, we used the established Shh gradient at t = 10 h as initial conditions and set S to 0 throughout the domain.

