## Supplementary Information for "Illuminating morphogen and patterning dynamics with optogenetic control of morphogen production"

Contains:

Supplementary Figures S1-S3

Supplementary Tables S1-S3

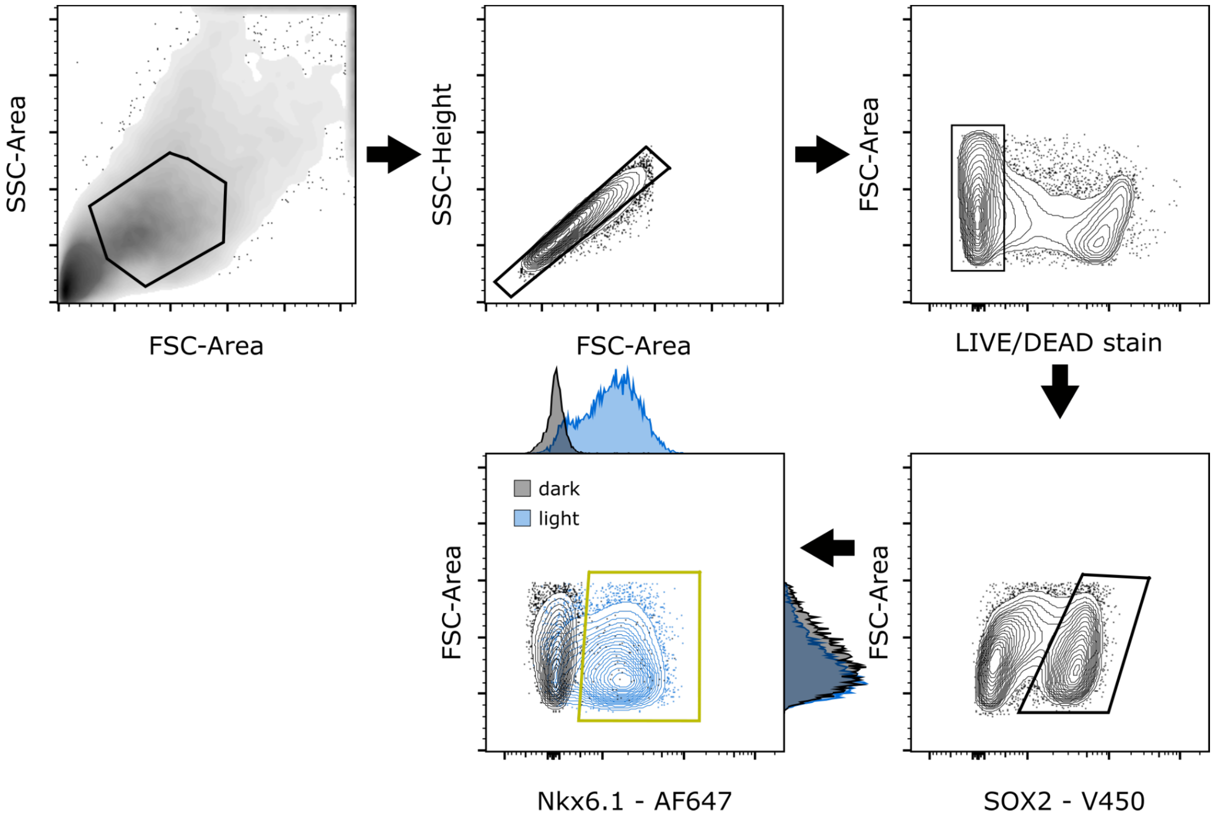


**Figure S1. Example of flow gating strategy.** Single cells were selected by forward and side scatter. Live cells were selected by gating for LIVE/DEAD stain negative cells. Neural progenitors were enriched by gating for Sox2-positive cells. Finally, the percentage of cells positive for a patterning marker was calculated. In this example, the effect of blue-light illumination on Nkx6.1 expression is illustrated for a cell line that expresses Shh under control of DD-GAVPO.

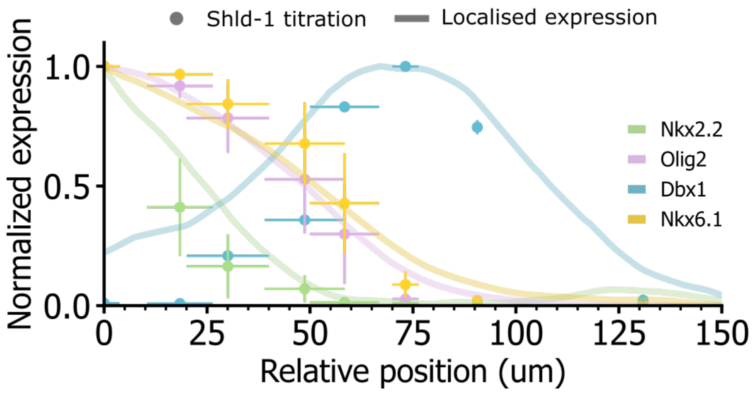


**Figure S2. Comparison between Shld-1 titration and spatial Shh gradient.** Using mScarlet fluorescence as a proxy for Shh expression, Shld-1 concentration dependent patterning gene expression (Fig. 1 H) was mapped onto an exponential gradient with a length-scale of 25 µM (mean and SEM of two independent experiments). This resulted in a good agreement with the patterning response to spatially localised Shh production (lines, distance from half-maximal mScarlet fluorescence), suggesting an underlying gradient with similar length-scale (data derived from Fig. 2).

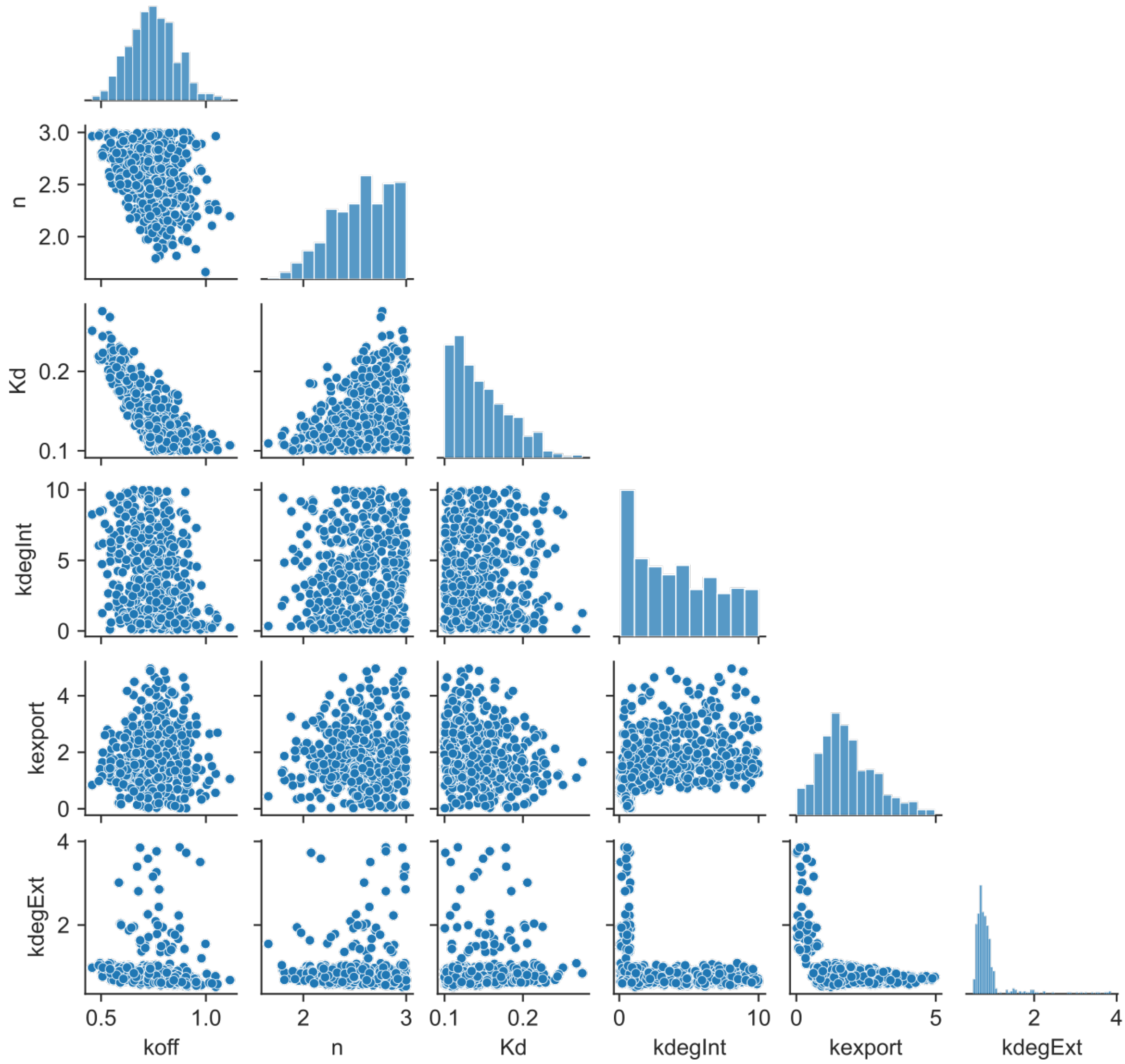


**Figure S3. Pairwise joint approximate posterior distributions for ABC-based parameter estimates.**

**Table S1.** Plasmids used for cell line construction. Promoters are represented by "pr", terminators are represented by "t".

| Plasmid | Backbone | Type | Insert | Source |
|---|---|---|---|---|
| PBgg-Puro | PB-U6insert | piggyBac vector with Puromycin resistance | - | this work |
| PBgg-Hygro | PB-U6insert | piggyBac vector with Hygromycin resistance | - | this work |
| pDBB9 | PBgg-Puro | piggyBac donor plasmid | prEF1α-GAVPO-tSV40 | this work |
| pDBB31 | PBgg-Puro | piggyBac donor plasmid | prEF1α-DD-GAVPO-tSV40 | this work |
| pDBB56 | PBgg-Hygro | piggyBac donor plasmid | prEF1α-Shh-IRES2-H2B-mScarletI-tSV40 | this work |
| pDBB57 | pJT371 | Cp36 donor plasmid | prEF1α-mScube2-tSV40 | this work |
| pDBB62 | PBgg-Hygro | piggyBac donor plasmid | prEF1α-NShh-IRES2-H2B-mScarletI-tSV40 | this work |
| pDBB63 | PBgg-Hygro | piggyBac donor plasmid | prEF1α-NShh-CD4-IRES2-H2B-mScarletI-tSV40 | this work |

**Table S2.** Mouse embryonic stem cell lines used in this study.

| Name | Insertions | Parental line | Data shown in Figure |
|---|---|---|---|
| DBO1 | prEF1α-Shh-IRES2-H2B-mScarletI-tSV40 (pDBB56) | HM1 | - |
| DBO2 | prEF1α-Shh-IRES2-H2B-mScarletI-tSV40 (pDBB56); prEF1α-GAVPO-tSV40 (pDBB9) | DBO1 | Fig. 1D-F |
| DBO3 | prEF1α-Shh-IRES2-H2B-mScarletI-tSV40 (pDBB56); prEF1α-DD-GAVPO-tSV40 (pDBB31) | DBO1 | Fig. 1D-F |
| DBO3-c4 (clonal) | prEF1α-Shh-IRES2-H2B-mScarletI-tSV40 (pDBB56); prEF1α-DD-GAVPO-tSV40 (pDBB31) | DBO3 | Fig. 1 G, H; Fig. 2; Fig. 3D; Fig. 4 |
| DBO4 | prEF1α-Shh-IRES2-H2B-mScarletI-tSV40 (pDBB56); prEF1α-DD-GAVPO-tSV40 (pDBB31), prEF1α-mScube2-tSV40 (pDBB57) | DBO3-c4 | Fig. 3C, D |
| DBO5 | prEF1α-DD-GAVPO-tSV40 (pDBB31) | HM1 | - |
| DBO6 | prEF1α-DD-GAVPO-tSV40 (pDBB31), prEF1α-Shh-IRES2-H2B-mScarletI-tSV40 (pDBB56) | DBO5 | Fig. 3A, B |
| DBO7 | prEF1α-DD-GAVPO-tSV40 (pDBB31), prEF1α-NShh-IRES2-H2B-mScarletI-tSV40 (pDBB62) | DBO5 | Fig. 3A, B |
| DBO8 | prEF1α-DD-GAVPO-tSV40 (pDBB31), prEF1α-NShh-CD4-IRES2-H2B-mScarletI-tSV40 (pDBB63) | DBO5 | Fig. 3A, B |

**Table S3.** Priors used for ABC-based parameter inference.

| Parameter | Description | Prior distribution |
|---|---|---|
| $k_{off}$ ($h^{-1}$) | GAVPO dark-state reversion rate | Uniform(0.1, 3.) |
| n (-) | hill coefficient | Uniform(1., 3.) |
| $K_d$ (-) | $GAVPO_{on}$ required for half-maximal Shh production | Uniform(0.1, 2.) |
| $k_{degInt}$ ($h^{-1}$) | degradation of cell-internal Shh | Uniform(0.1, 10.) |
| $k_{export}$ ($h^{-1}$) | rate of Shh export | Uniform(0.01, 10.) |
| $k_{degExt}$ ($h^{-1}$) | degradation of cell-external Shh | Uniform(0.1, 4.) |
